# The effect of subthreshold transcranial magnetic stimulation on the excitation of corticospinal volleys with different conduction times

**DOI:** 10.1101/084574

**Authors:** Niclas Niemann, Patrick Wiegel, John C. Rothwell, Christian Leukel

## Abstract

Previous studies in humans interested in inhibitory synaptic activity at the level of the primary motor cortex (M1) have frequently used an electrophysiological technique call short intracortical inhibition (SICI). This technique consists of two subsequent pulses with transcranial magnetic stimulation (TMS), of which the first pulse (S1) has been argued to target local inhibitory connections, reducing corticospinal output generated by the second pulse (S2). However, the reduction of corticospinal output is not seen in every tested subject, sometimes S1 even increased corticospinal output. Thus there seems to be more occurring than just the targeting of local inhibitory connections by S1, indicating that the mechanisms of SICI are not fully understood. In the present study, in 18 healthy young subjects we applied a method allowing to segregate corticospinal volleys with different conduction times and investigated the effect of S1 on the excitation of these volleys. Our results revealed three major findings, which can be summarized as follows: i) S1 acted not only locally at the level of M1, but produced corticospinal activity which may influence corticospinal output by S2 at the spinal level; ii) there was not only reduced excitation of corticospinal volleys by S1, but also an increased excitation of corticospinal volleys with longer conduction times; iii) at the level of M1, S1 indeed targeted preferentially inhibitory synaptic connections, and these reduced corticospinal excitation of the fastest conduction corticospinal volleys.

Our results indicate that underlying mechanisms in studies applying SICI should be interpreted with caution.

## Introduction

Inhibition plays a crucial role for the functioning of the nervous system (Isaacson and Scanziani 2011; Letzkus et al. 2015). In studies on humans, there exists an electrophysiological technique called short interval intracortical inhibition (SICI) which is considered to probe local activity of synaptic inhibition in the primary motor cortex (M1). This method was developed over 20 years ago (Kujirai et al. 1993) and has since then been frequently applied to test modulations in the activity of synaptic inhibition with respect to movement control (e.g. Barthelemy and Nielsen 2010) and motor learning (e.g. Perez et al. 2004) in the context of degeneration (e.g. Heise et al. 2013) and disease (e.g. Beck et al. 2008).

SICI is based on the application of two subsequent pulses with transcranial magnetic stimulation (TMS) to the primary motor cortex (M1). The stimulation intensity of the two pulses is different, the first stimulus (S1) is applied below the threshold for evoking a motor evoked potential (MEP) in muscles of the contralateral side, and the second stimulus (S2) is applied at an intensity at or above the threshold for evoking an MEP. The MEP in the target muscle produced by S2 was lower in size when S1 and S2 were combined than when S2 was applied without S1. Pharmacological studies provided evidence that the mechanism of this reduction is related to GABAAergic synapses that are preferentially activated by TMS below MEP threshold (Muller-Dahlhaus et al. 2008; Ziemann et al. 2015). The fact that S1 produced no MEP has led to the believe that there is no corticospinal activity by S1. Thus the reduction of the MEP by S1 has been attributed to activation of local GABAAergic inhibitory connections in M1 that reduce the corticospinal output produced with S2 (Rothwell et al. 2009). However, SICI cannot be displayed in every subject. In about 20% of the cases, there is no inhibition, but rather no effect or even an increase of the MEP by S1 (Wassermann 2002). Especially the large increase of the MEP seen in some subjects is problematic, as it contradicts with the postulated mechanism of SICI, namely the recruitment of local inhibitory intracortical connections with TMS applied below MEP threshold. Given the great importance of a tool that enables to study inhibitory synaptic activity in the human cortex it seems worthwhile to better understand the mechanisms of SICI. Therefore, in the present experiment, we studied the effect of S1 on the excitation of corticospinal volleys with different conduction times using an electrophysiological technique called H-reflex conditioning (Nielsen et al. 1993). The MEP reflects the summation of activity of all corticospinal neurons and connections that are activated by the pulse. H-reflex conditioning allows to segregate the corticospinal volleys pooled in the MEP, which arrive at the spinal motoneurons with different time delays with respect to the application of the TMS pulse. Thus the aim of the present experiment was to assess how S1 influences the excitation of corticospinal volleys produced by the subsequent S2.

## Methods

### Subjects

18 healthy subjects (aged 25 ± 3 years (mean ± standard deviation, STD), 8 women) participated in the study. Exclusion criterions for participation were metal implants in eyes and head, pregnancy and any neurological disease (Rossi et al. 2009). Each subject participated in a single experimental session. Prior to the experiment, all participants gave written informed consent to the procedures, which were in accordance with the latest revision of the Declaration of Helsinki (Fortaleza, Brazil, 2013) and were approved by the local ethics committee of the Albert-Ludwigs-Universitöt in Freiburg (423/15). Subjects were compensated for participation in form of a small monetary reward.

### Electromyography (EMG)

Surface EMG was recorded from the left soleus (SOL) and tibialis anterior (TA) muscles using bipolar surface electrodes (Blue sensor P, Ambu®, Bad Nauheim, Germany). For preparation, the skin was shaved, abraded with fine-grain sandpaper, and cleaned with isopropyl(70%)/water(30%) solution. Electrodes were attached on the muscle belly with 2 cm interelectrode distance, and a ground electrode was placed on the tibial plateau. Impedance was below 10 kΩ. EMG signals were pre-amplified in the sensors (TA × 500; SOL × 100), further amplified (2 ×), bandpass filtered (10 – 1000 Hz) and sampled at 2 kHz. All data was stored for later offline analysis using a custom-built software (www.pfitec.de; LabView based; National Instruments®, Austin, TX, USA).

### Electrophysiological stimulation techniques

All measurements were performed with the subjects at rest. Subjects were seated comfortably in a custom-built laboratory seat with a neck support. We asked the subjects to keep their head in a stable position and their legs placed on a custom built footboard in a stretched but relaxed position (Figure 1 A).

#### Peripheral nerve stimulation (PNS)

SOL H-reflexes were elicited with a constant current stimulator (DS7a, Digitimer®, Hertfordshire, UK) by stimulating the posterior tibial nerve at the popliteal fossa. Stimuli consisted of square-wave pulses of 0.5 ms duration. A graphite coated rubber pad of 5 × 5 cm was used as anode and was fixed on the anterior aspect of the knee just underneath the patella. The cathode (displaceable custom built 1.5 cm diameter AgCL electrode) was placed in the popliteal fossa and changed stepwise until the optimum site for eliciting the SOL H-reflex was found (stimulation frequency of 4 s). The optimum site was defined as the site where low stimulation intensity (in between 5 and 20 mA) elicited a consistent SOL H-reflex with no or a small SOL M-wave. Further, stimulation at this optimum site did not activate the common peroneal nerve, which was tested with parallel recordings from the TA muscle (TA H-reflex and TA M-wave). After the optimum site was found, a self-adhesive cathode (Blue sensor P, Ambu®, Bad Nauheim, Germany) was fixed at this site. We recorded H/M recruitment curves from electrical stimuli applied every 4 s. First we delivered a few stimuli to pinpoint in which range of intensities the maximum H-reflex (Hmax) and the maximum M-wave (Mmax) could be evoked. Thereafter we applied approximately 40-50 stimuli in smaller intensity steps around Hmax, Mmax, and the upsloping part of the H-reflex curve. We determined Hmax and Mmax from these recordings for each subject. Hmax and Mmax amplitudes were used for calculating the intensity of PNS for the conditioned H-reflex measurement (see paragraph *“Conditioned H-reflexes”*). The stimulation intensity for PNS for conditioned H-reflexes was set to a fixed value, which consistently produced H-reflexes of a size that matched 15 to 25% of the individual Mmax. The criterion of 15 – 25% Mmax has been argued to ensure that the output of the spinal motoneurons to a conditioning input is linear and that the spinal motoneurons are equally sensitive to both excitatory and inhibitory input (Crone et al. 1990). The grand mean value for the H-reflex size during conditioned H-reflex measurement was 19 ± 6% of Mmax (mean ± STD) recorded before the H-reflex conditioning measurement in the present study, which means that the mentioned criterion was satisfied. H/M recruitment curves were recorded again after the conditioned H-reflex measurement to identify neural modulations at the spinal and peripheral level in the time course of the experiment.

#### Transcranial magnetic stimulation (TMS)

Single pulse and paired pulse TMS was applied over the contralateral primary motor cortex of the leg area (M1), using a Magstim® 200^2^ Stimulator with a BiStim unit (Magstim® Company Ltd., Whitland, UK) and a figure-of-eight batwing coil (70mm) or a double cone coil (90mm). The double cone coil was used if the stimulation intensity required to produce motor evoked potentials (MEPs) in SOL could not be reached with the batwing coil. The coil was positioned on the scalp with a 0° angle in the sagittal plane. The current flow was from posterior to anterior. We conducted a mapping procedure to identify the optimal site on the scalp for eliciting SOL MEPs (SOL hotspot). The optimum site was defined as the site where MEPs could be clearly evoked with the lowest possible stimulation intensity. Resting motor threshold (RMT) was determined for TMS at the SOL hotspot. RMT was determined as the lowest intensity of the magnetic stimulator that was required to evoke MEPs of 50 μV peak-to-peak amplitude in at least three out of five consecutive trials (Kujirai et al. 1993; Rossini et al. 1994). The grand mean stimulation intensity at RMT was 54 ± 7% (mean ± STD) of the maximum stimulator output. To ensure a constant location of the coil throughout the experiment relative to the subject’s head, the handle of the coil was fixed by a mounting tool (Manfrotto® Magic Arm, Lino Manfrotto & Co, Cassola, Italy) that was fixated at the headrest of the seat. The position of the subjects’ head was stabilized by a custom-built neck brace mounted to the headrest. The coil position was marked on the scalp with a waterproof pen and monitored throughout the experiment by the experimenter.

#### Conditioned H-reflexes

H-reflex conditioning with TMS to M1 was already applied in previous experiments (Nielsen et al. 1993; Nielsen et al. 1995; Taube et al. 2011; Leukel et al. 2012; Leukel et al. 2015; Taube et al. 2015).

The H-reflex conditioning technique takes advantage of the fact that different corticospinal volleys induced by TMS have different conduction times and thus will arrive at spinal motoneurons with different latencies. The objective of this technique is to promote a coincidence of TMS evoked corticospinal volleys and an afferent volley by PNS at the spinal motoneurons. To provoke a collision between distinct corticospinal volleys and the afferent volley, PNS relative to TMS is applied with different delays, termed interstimulus intervals (ISIs). Negative ISIs indicate that the application of PNS precedes the application of TMS and positive ISIs indicate the opposite.

PNS without the additional application of TMS produces an H-reflex. The combination of the two methods PNS and TMS produces a conditioned H-reflex, as the corticospinal volley changes the recruitment of spinal motoneurons when compared to the excitation of spinal motoneurons by afferents alone. The fastest corticospinal volley originating from corticomotoneuronal cells in M1 (Lemon 2008) excite spinal motoneurons faster than afferent fibres by PNS. The “first increase” of the conditioned H-reflex size when testing from negative to positive ISIs is termed “early facilitation”, and results from the additional (besides PNS) excitation of spinal motoneurons by corticomotoneuronal cells (Nielsen et al. 1993). The early facilitation is usually seen at ISIs in the range between -4 ms and -1 ms. This inter-individual difference of 3 ms is caused by differences in physiological and anatomical parameters like leg length and trunk length.

As mentioned in paragraph “*Peripheral nerve stimulation*” the intensity of PNS for evoking H-reflexes was set to the defined criterion (15–25% of the individual Mmax) for each subject. The intensity of TMS was set to 1.2 × RMT to ensure a consistent corticospinal excitation produced by the stimulation. The grand mean value of the stimulation intensity at 1.2 RMT was 64 ± 9% (mean ± STD) of the maximum stimulator output. PNS and TMS were applied with 14 different ISIs ranging from -5 ms to +8 ms, in steps of 1 ms.

#### Short interval intracortical inhibition (SICI)

SICI refers to an established method of two subsequent TMS pulses, applied with a single delay in the range of 1 ms to 5 ms (Kujirai et al. 1993). SICI in our study was combined with the H-reflex conditioning technique. This means to include a second, subthreshold TMS pulse (S1) preceding the suprathreshold TMS pulse (S2) used for H-reflex conditioning.

Two distinct delays between these two TMS pulses were tested, resulting in two SICI conditions with 1 ms (Hcond_SICI_1_) and 2.5 ms (Hcond_SICI_2.5_) between S1 and S2, respectively. The two intervals were argued to produce different types of cortical inhibitory effects (Fisher et al. 2002; Chen 2004).

The intensity of the conditioning S1 pulse was determined by a testing procedure that was performed before conducting the H-reflex conditioning measurement. This test procedure consisted of several blocks of trials. In each block, S2 alone and the combination of S1 and S2 with a delay of 2.5 ms (SICI_2.5_) were applied in a randomized order. Twenty trials (10 for S2 alone, 10 for the combination of S1 and S2) were recorded in each block. The pause between successive trials was 4 s. Stimulation intensity for S2 was 1.2 RMT, and the stimulation intensity for S1 was varied in-between blocks, from 0.45 RMT to 0.75 RMT in steps ranging from 0.1 to 0.5 RMT. The objective of this testing procedure was to find the highest decreasing effect of S1 on the SOL MEP size produced by S2. The stimulation intensity of S1 producing the maximum reduction of the S2 MEP was used for the H-reflex conditioning measurement. The grand mean value of this stimulation intensity for S1 was 0.66 ± 0.04 RMT (mean ± STD), corresponding to a stimulator output of 35 ± 5% of the maximum stimulator output (mean ± STD).

### H-reflex conditioning + SICI measurement

All three stimulation conditions (Hcond; Hcond_SICI_2.5_; Hcond_SICI_1_) were tested at once in a large stimulation protocol (Figure 1B). We decided to not test the different conditions in a sequential order, as effects produced by one condition may have influenced effects produced by the subsequent condition.

The various variables contained in these conditions (Hcond; Hcond_SICI_2.5_; Hcond_SICI_1_) and several control variables were recorded. There were 48 different variables, and each of these variables was tested in a randomized order:

- 14 × Hcond (14 ISIs, ranging from -5 to +8 ms): assessing the size of conditioned H-reflexes
- 14 × Hcond SICI_2.5_ (14 ISIs, ranging from -5 to +8 ms, ISIs according to the delay between PNS and TMS pulse S2): assessing the size of conditioned H-reflexes)
- 14 × Hcond SICI_1_ (14 ISIs, ranging from -5 to +8 ms, ISIs according to the delay between PNS and TMS pulse S2): assessing the size of conditioned H-reflexes
- 1 × PNS without TMS: assessing the size of the unconditioned H-reflex
- 1 × TMS S2 without PNS: assessing the size of the MEP (MEP S2)
- 1 × TMS S1 without PNS: assessing the size of the MEP in case S1 would produce muscular activity (MEP S1)
- 1 × SICI_2.5_ without PNS: assessing the size of the SICI MEP (MEP_SICI_2.5_)
- 1 × SICI_1_ without PNS: assessing the size of the SICI MEP (MEP_SICI_1_)
- 1 × PNS combined with the just the TMS pulse S1 without S2, at an ISI of +1 ms: assessing the size of the H-reflex that is conditioned by S1 (Conditioned H-reflex_S1)

After each of the 48 variables was recorded in one cycle of stimuli, a new set of 48 trials started. Subjects completed a total of 15 cycles of 48 trials. Thus a total number of 720 trials were recorded in an experiment. The pause between trials was always 4 s. The reason for keeping the pause constant relates to a potential bias of the results by changes in post activation depression acting on Ia afferent fibres at the spinal level (Crone and Nielsen 1989). Modulations in the strength of post activation depression by variations of the delay between repeated PNS application drastically changes the size of the H-reflex size (Crone and Nielsen 1989). Thus interpretations about changes in corticospinal excitation deduced from conditioned H-reflexes could be biased in case of changes in post activation depression.

**Figure 1.**
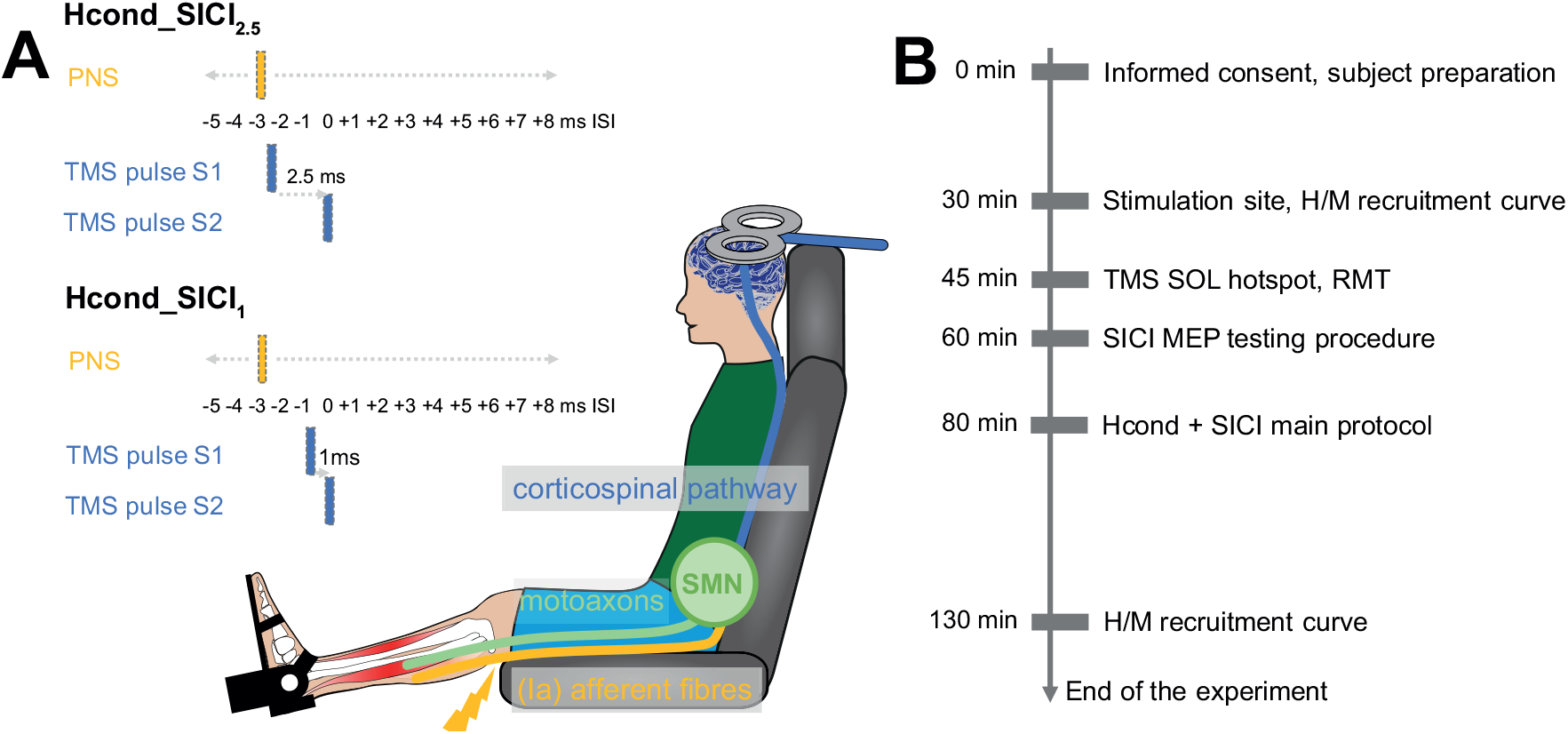
**A** depicts the experimental setup of H-reflex conditioning in combination with SICI. The coloured boxes on the left side indicate the instants of the stimulus/pulses for PNS/TMS. In the example depicted in the figure, the timing of the stimulus/pulses reflects the testing of ISI -3 ms, both for Hcond_SICI_2.5_ and Hcond_SICI_1_. Note that the delay between S1 and S2 remained constant whereas the delay between PNS and S1/S2 was varied when testing ISIs from -5 ms and +8 ms, in steps of 1 ms. SMN: spinal motoneurons. The figure was adapted from Lundbye-Jensen et al. (2011). **B** shows the experimental protocol. The timeline on the left side of the horizontal grey bars refer to an experiment running under ideal conditions. The time to complete one experiment therefore reached from 130 min to 170 min.

### Data analysis

Peak-to-peak amplitudes of the unrectified SOL EMG were calculated for all MEPs, conditioned and unconditioned H-reflexes as well as for maximum H-reflex and maximum M-waves. The results of trials corresponding to the different variables were averaged for each individual.

The first increase in size of conditioned H-reflexes from negative to positive ISIs is termed “early facilitation” (see paragraph *“Conditioned H-reflexes*”). To identify the early facilitation in each individual, we first looked at the mean plots of conditioned H-reflexes from the Hcond condition and judged at which ISI, from -5 ms, to -4 ms, to -3 ms, etc., there was a prominent increase in the size of the mean conditioned H-reflexes compared to mean conditioned H-reflexes at the more negative ISIs. This is typically very easy to identify, as there occurs a visible increase in almost every tested subject between ISI -4 ms and ISI -1 ms. To test whether this increase was statistically significant, we performed uncorrected paired Student’s t-tests (P < 0.05) between conditioned H-reflexes at the ISI that showed the prominent increase and conditioned H-reflexes at the ISI with a time lag of -1 ms with respect to this ISI. We further performed uncorrected paired Student’s t-tests between conditioned H-reflexes at the ISIs with a time lag of -1 ms and the ISI -2 ms with respect to the ISI showing the prominent increase, and the same comparison for ISIs -2 ms and -3 ms. In all but three subjects, there was a significant difference in the size of conditioned H-reflexes between the ISI showing the prominent increase and the ISI with a time lag of -1 ms relative to this ISI, but no significant difference for comparisons between ISIs with more negative time lags. In the three subjects that did not show a statistical difference, we nevertheless denoted the ISI that we selected visually as early facilitation. Note that, in case the prominent increase in a subject occurred at ISI -4 ms, we performed paired Student’s t-tests between ISI -4 ms and ISI -5 ms, and between ISI -5 ms and the unconditioned H-reflexes, to ensure that the latter comparison did not reveal significant results. Four subjects showed the early facilitation at ISI -4 ms, 12 subjects showed the early facilitation at ISI -3 ms, and 2 subjects showed the early facilitation at ISI -2 ms.

The next step was to reference the mean of the conditioned H-reflexes at each ISI to the mean of the unconditioned H-reflex in every subject. We decided to consider not only the recorded unconditioned H-reflexes as reference, but also conditioned H-reflexes recorded at more negative ISIs than the early facilitation. The rationale for also including more negative ISIs than the early facilitation is that spinal motoneurons are expectedly not influenced by corticospinal input at these ISIs. They can therefore serve as reference H-reflexes and by reducing the natural physiological variation of the H-reflex – because more trials can be averaged – improve the reliability of the reference. Importantly, when we analysed the data we found out that only data from the Hcond condition should be included for calculating the reference H-reflex, but not data from the Hcond SICI_2.5_ and Hcond SICI_1_ conditions. We will refer to this issue in detail in the “Results” and “Discussion” section. We therefore included conditioned H-reflexes at more negative ISIs only from the Hcond condition for calculating the reference H-reflex. The intra-individual mean of the conditioned H-reflex obtained at each ISI, for all tested conditions (Hcond, Hcond SICI_2.5_ and Hcond SICI_1_) was divided by the intra-individual reference H-reflex and then multiplied by 100 to obtain the percentage of response with respect to the reference value. Finally, referenced conditioned H-reflexes were aligned to the ISI of the individual early facilitation. Thus the ISIs in the “Results” section refer to this alignment, and are thus denoted as EFD (delay with respect to the early facilitation, e.g. EFD -1ms; EFD +1 ms).

### Statistical analysis

All data sets showed normality and homogeneity, which was tested by the Kolmogorov-Smirnov test and the Levene’s test, respectively.

We calculated a power analysis according to the assumptions made by Faul et al. (2007) to assess whether the number of analysed subjects was sufficient to reveal statistical differences. A moderate effect size of 0.32, a statistical power of 80% and a ɑ-level of 0.05 was assumed. The result of this analysis showed that 18 subjects are sufficient (F-test; repeated-measures ANOVA, within factors; one group; three measurements).

#### Differences of referenced conditioned H-reflexes between conditions

To assess differences of referenced conditioned H-reflexes between the stimulation conditions, a repeated-measures ANOVA (3 × 12) with the factors CONDITION (Hcond, Hcond_SICI_2.5_, Hcond_SICI_1_), and EFD (-2 ms, -1 ms, 0, +1, +2, +3, +4, +5, +6, +7, +8, +9, +10 ms and +11 ms) was performed. Note that this selection of the range of EFDs is associated with few missing values at EFD -2 ms. The alignment to the individual early facilitation caused these missing values. Four subjects in our experiment showed the early facilitation at ISI -4 ms. As we applied conditioned H-reflexes in the range from ISIs -5 ms to +8 ms, referenced conditioned H-reflexes at EFD -2 ms were missing in these four subjects.

In addition to the ANOVA, post hoc paired Student’s t-tests were performed at each EFD from -2 ms to +11 ms between the different conditions (Hcond and Hcond_SICI_2.5_, Hcond and Hcond_SICI_1_, Hcond_SICI_2.5_ and Hcond_SICI_1_). Results obtained from these multiple comparisons were corrected by the Benjamini-Hochberg procedure (Benjamini and Hochberg 1995).

#### Differences of Hmax, Mmax, and H/M-ratio

To test for possible neural modulations at the spinal and peripheral level throughout the experiment, the variables Hmax, Mmax as well as the H/M-ratio from pre-and post-tests were analysed with paired Student’s t-tests.

*Control variables: MEP S2, MEP S1, MEP_SICI_2.5_, MEP_SICI_1_, Unconditioned H-reflex, Conditioned H-reflex_S1*

An ANOVA with the factor CONDITION was used for the comparison of MEPs (*MEP S2, MEP_SICI_2.5_, MEP_SICI_1_*). Paired Student’s t-tests were performed for further comparisons: MEP S2 and MEP_SICI_2.5_; MEP S2 and MEP_SICI_1_; MEP_SICI_2.5_ and MEP_SICI_1_; MEP S2 and MEP S1; unconditioned H-reflex and conditioned H-reflex_S1. t-tests for MEPs were corrected by the Benjamini-Hochberg procedure (Benjamini and Hochberg 1995).

The level of significance was set to *P* < 0.05 for all tests. Mean values and standard error of the mean (SEM) are reported unless indicated otherwise. Greenhouse-Geisser corrected values for ANOVAs are reported in case sphericity of the tested samples was violated. Data were statistically analysed with SPSS software 22.0 (SPSS®, Chicago, IL, USA).

## Results

### Referenced conditioned H-reflexes

The referenced conditioned H-reflex grand mean curves for each condition (Hcond, Hcond_SICI_1_, Hcond_SICI_2.5_) are depicted in Figure 2A.

The ANOVA yielded no effect for CONDITION (*F*_2,34_ = 0.857, *P* = 0.0433), a significant effect for EFD (*F*_11,187_ = 8.593, *P* < 0.001) and a significant effect for CONDITION × EFD (*F*_22,374_ = 12.605, *P* < 0.001). This result confirms the visual impression of the shape of the mean curves displayed in Figure 2 A. The size of referenced conditioned H-reflex in all conditions was different between EFDs (main effect of EFD). The time course of facilitation and depression of referenced conditioned H-reflexes was different between conditions (interaction CONDITION × EFD).

Differences between the three conditions at each EFD were evaluated with paired Student’s t-tests (Table 1 A). These results can be summarized as follows:

a. Compared with the control condition (Hcond), Hcond_SICI_2.5_ and Hcond_SICI_1_ both demonstrated less H-reflex facilitation for EFD 0 ms and EFDs +3 to +5 ms whilst they showed more facilitation for EFDs +8 to +10 ms. There was no difference between the conditions at EFDs +6, +7 and +11 ms.
b. For EFD 0 ms and EFDs +3 to +11 ms, there was no difference between Hcond_SICI_2.5_ and Hcond_SICI_1_. Hcond_SICI_2.5_ and Hcond_SICI_1_ differed at EFDs -2, -1, +1 and +2 ms. At EFD -2 ms, Hcond_SICI_2.5_ facilitated referenced conditioned H-reflexes whereas it depressed them at EFD -1 ms.
c. At EFD +1 ms, Hcond_SICI_2.5_ showed no different facilitation as the control condition (Hcond) while the referenced conditioned H-reflex in the Hcond_SICI_1_ condition was depressed. Conversely at EFD +2 ms, Hcond_SICI_1_ was not different from Hcond while referenced conditioned H-reflexes in the Hcond_SICI_2.5_ condition was depressed.

**Table 1.** **A** shows the P-values from the paired Student’s t-tests for the comparison of referenced conditioned H-reflexes at EFDs from -2 ms to +11 ms between the different conditions. The results of Student’s t-tests were adjusted for multiple comparisons by the Benjamini and Hochberg procedure (Benjamini and Hochberg 1995). The corrected significance level (P-value) is displayed in the right column. Comparisons that reached below this significance level are marked in green colour. **B** shows the P-values from the paired Student’s t-tests that were calculated for the reduced dataset of 10 subjects. For these analyses, we discarded subjects that showed a significant difference of referenced conditioned H-reflexes at EFD -2 ms or EFD -1 ms between the conditions Hcond and Hcond_SICI_2.5_. This was the case in 4 subjects. Four other subjects had to be excluded from the analyses as they had missing values at EFD -2 ms. The grey coloured columns refer to EFDs where the three comparisons showed the same results for the full dataset (A) and the reduced dataset (B).

|  | -2 | -1 | 0 | +1 | +2 | +3 | +4 | +5 | +6 | +7 | +8 | +9 | +10 | +11 | correction |
| --- | --- | --- | --- | --- | --- | --- | --- | --- | --- | --- | --- | --- | --- | --- | --- |
| **A** Hcond and Hcond_SICI<sub>2.5</sub> | 0.02 | 0.03 | 0.003 | 0.94 | 0.01 | 0.01 | <0.001 | 0.002 | 0.05 | 0.74 | <0.001 | <0.001 | 0.02 | 0.07 | <0.036 |
| Hcond and Hcond_SICI<sub>1</sub> | 0.51 | 0.032 | 0.001 | 0.03 | 0.34 | 0.01 | <0.001 | 0.01 | 0.38 | 0.37 | <0.001 | <0.001 | 0.024 | 0.33 | <0.032 |
| Hcond_SICI<sub>2.5</sub> and Hcond_SICI<sub>1</sub> | 0.01 | 0.002 | 0.8 | 0.002 | 0.004 | 0.75 | 0.31 | 0.1 | 0.08 | 0.14 | 0.39 | 0.21 | 0.48 | 0.12 | <0.014 |
| **B** Hcond and Hcond_SICI<sub>2.5</sub> | 0.08 | 0.34 | <0.001 | 0.28 | 0.023 | 0.03 | <0.001 | 0.006 | 0.008 | 0.83 | 0.004 | 0.001 | 0.1 | 0.11 | <0.025 |
| Hcond and Hcond_SICI<sub>1</sub> | 0.28 | 0.09 | 0.0137 | 0.143 | 0.5 | 0.03 | <0.001 | 0.05 | 0.24 | 0.40 | 0.004 | 0.004 | 0.08 | 0.23 | <0.0142 |
| Hcond_SICI<sub>2.5</sub> and Hcond_SICI<sub>1</sub> | 0.02 | 0.04 | 0.85 | 0.03 | 0.02 | 0.56 | 0.26 | 0.002 | 0.22 | 0.44 | 0.47 | 0.8 | 0.93 | 0.34 | <0.003 |

The facilitation of referenced conditioned H-reflexes at EFD -2 ms in the Hcond_SICI_2.5_ condition was unexpected since this is at a time interval when, by definition, there is no modulation of the H-reflex. The modulation starts at EFD 0 ms. As we discuss, the most likely reason is that S1 itself produced a small descending corticospinal volley. This highly likely accounts for the H-reflex facilitation observed at -2 ms.

The intensity of S1 was adjusted on an individual basis, producing clear depression of the referenced conditioned H-reflex at EFD 0 ms. This S1 intensity ranged from about 0.6 – 0.7 RMT between individuals. We therefore tested whether the degree of facilitation of the referenced conditioned H-reflex at EFD -2 ms was related to the S1 intensity expressed as a fraction of RMT. Fig 3 B shows that there was no correlation between these variables (r = 0.34, P = 0.16).

Based on the postulated mechanism of SICI, we expected that S1 will depress the response to S2 via local inhibitory interactions in the cortex. If so we reasoned that the excitatory effect at -2 ms should have a different mechanism than the depression at EFD 0 ms. We therefore examined whether there was a correlation between the effects of Hcond_SICI_2.5_ at EFD -2 ms and 0 ms. Fig 3 A shows that there was no significant relationship, consistent with the existence of separate mechanisms (r = 0.35, P = 0.22). Further, as there was a significant depression of referenced conditioned H-reflexes at EFD -1 ms with Hcond_SICI_2.5_ besides the facilitation at EFD -2 ms, we examined whether there was a correlation between the effects at EFD -1 ms and 0 ms. Figure 3 A shows that there was no such relationship (r = 0.28, P = 0.23), also indicating separate underlying mechanisms.

To confirm these findings, in further analyses, we discarded participants that showed a significant facilitation of the referenced conditioned H-reflexes by Hcond_SICI_2.5_ at EFD -2 ms or a significant depression at EFD -1 ms. Therefore, we calculated paired Student’s T-tests for the 15 recorded trials between Hcond_SICI_2.5_ and Hcond in each subject. The subject was discarded in case the P-value dropped below 0.05 at one of the two EFDs. This was the case in 4 subjects (Figure 3C), and the data from the remaining 10 subjects is depicted in Figure 2 B. Like for the full data set, we calculated paired Student’s t-tests for referenced conditioned H-reflexes at each EFD between the conditions, and the results are displayed in Table 1 B. Although there was no longer a significant difference of referenced conditioned H-reflexes between the conditions Hcond and Hcond_SICI_2.5_ at EFDs -2 and -1 ms, the reduction of conditioned H-reflexes at EFD 0 ms and also at EFD 4 ms by Hcond_SICI_2.5_ remained (see Table 1 B and Figure 3 C). Further, there was still a significant increase of conditioned H-reflexes with Hcond_SICI_2.5_ at EFDs +8 and +9 ms compared to Hcond. These findings suggest that the reduction of corticospinal excitation from S2 at least for some corticospinal volleys was not caused by descending corticospinal activity induced by S1.

**Figure 2.**
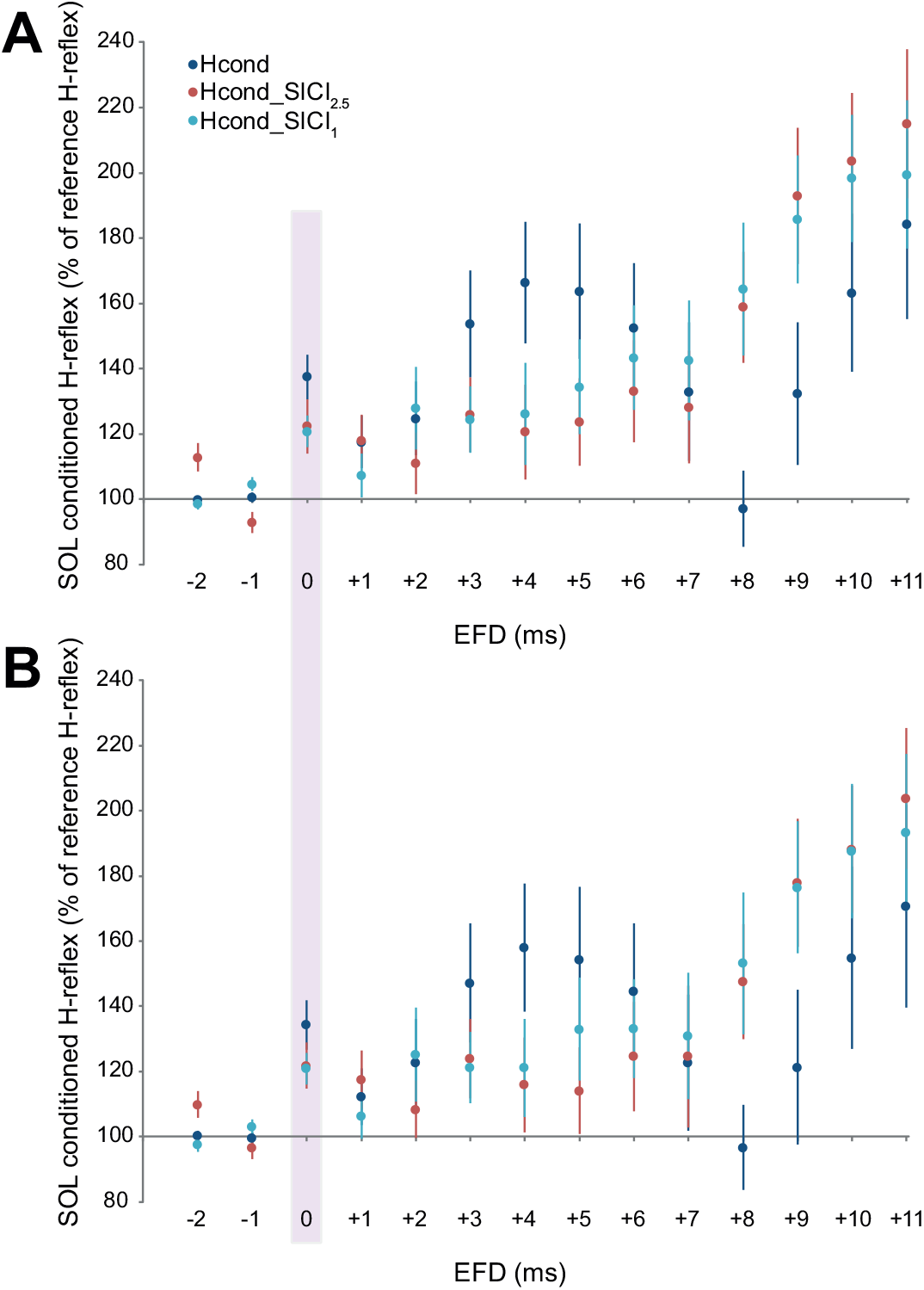
**A** shows referenced conditioned H-reflexes (mean ± SEM) for all conditions. **B** depicts the same data but for the reduced dataset of 10 subjects. Eight subjects were discarded because they showed significant differences of referenced conditioned H-reflexes at EFD -2 ms or EFD -1 ms between the conditions Hcond and Hcond_SICI_2.5_ (4 subjects), or had missing values at EFD -2 ms (4 subjects). The purple shaded area highlights the early facilitation (EFD 0 ms).

### Hmax, Mmax, H/M-ratio

Mean values of Hmax, Mmax and the H/M-ratio for pre-and post-test measurements are displayed in Figure 3 F.

Paired Student’s t-tests revealed no significant differences between the pre-and post-tests for Hmax (Hmax: *P* = 0.73), Mmax (Mmax: *P* = 0.56) and the H/M-ratio (H/M-ratio: *P* = 0.15). The results for Hmax indicate that the maximum excitation of spinal motoneurons by the afferent input did not change between the beginning and the end of the experiment. Unchanged Mmax values indicate that the recording conditions did not change between pre-and post-tests.

### Control variables

In these experiments, the intensity of S2 was 1.2 RMT. Thus S2 evoked an MEP which we could measure in trials without PNS. We could therefore test whether this MEP was depressed by S1 as expected by SICI; we could also test whether S1 itself was subthreshold for evoking an MEP at rest. The means of the amplitudes of MEP S2, MEP S1, MEP_SICI_2.5_, MEP_SICI_1_, unconditioned H-reflex, Conditioned H-reflex_S1 are displayed in Figure 3 D and Figure 3 E. For the MEPs (MEP S2, MEP_SICI_2.5_, MEP_SICI_1_), the ANOVA yielded a significant effect for CONDITION (*F*_2,50_ = 40.5, *P* < 0.001). As expected from SICI, paired Student’s t-tests showed that MEP_SICI_2.5_ and MEP_SICI_1_ were both smaller than MEP_S2 (*P* < 0.01 for both comparisons) but there was no difference between MEP_SICI_2.5_ and MEP_SICI_1_ (*P* = 0.171) (Figure 3E). The level of significance had to be lowered to 0.03 according to Benjamini and Hochberg (1995).

Student’s t-test revealed MEP_S2 was larger than MEP_S1 (*P* < 0.01) (Figure 3E); indeed, there was no response in the EMG detectable after S1.

For the comparison between the unconditioned H-reflex and the H-reflex conditioned with S1, Student’s t-test revealed no difference between the unconditioned H-reflex and the conditioned H-reflex_S1 (*P* = 0.454) (Figure 3D). This result indicates that the subthreshold TMS pulse S1 did not modulate the H-reflex at least for the chosen delay between PNS and S1 of +1 ms. Note that the ISI of +1 ms refers to an absolute ISI, not aligned to the early facilitation. With the early facilitation occurring at ISIs -4 ms, -3 ms or -2 ms, the ISI of +1 ms reflects the ISIs +5 ms, +4 ms or +3 ms, respectively, with respect to the early facilitation.

**Figure 3.**
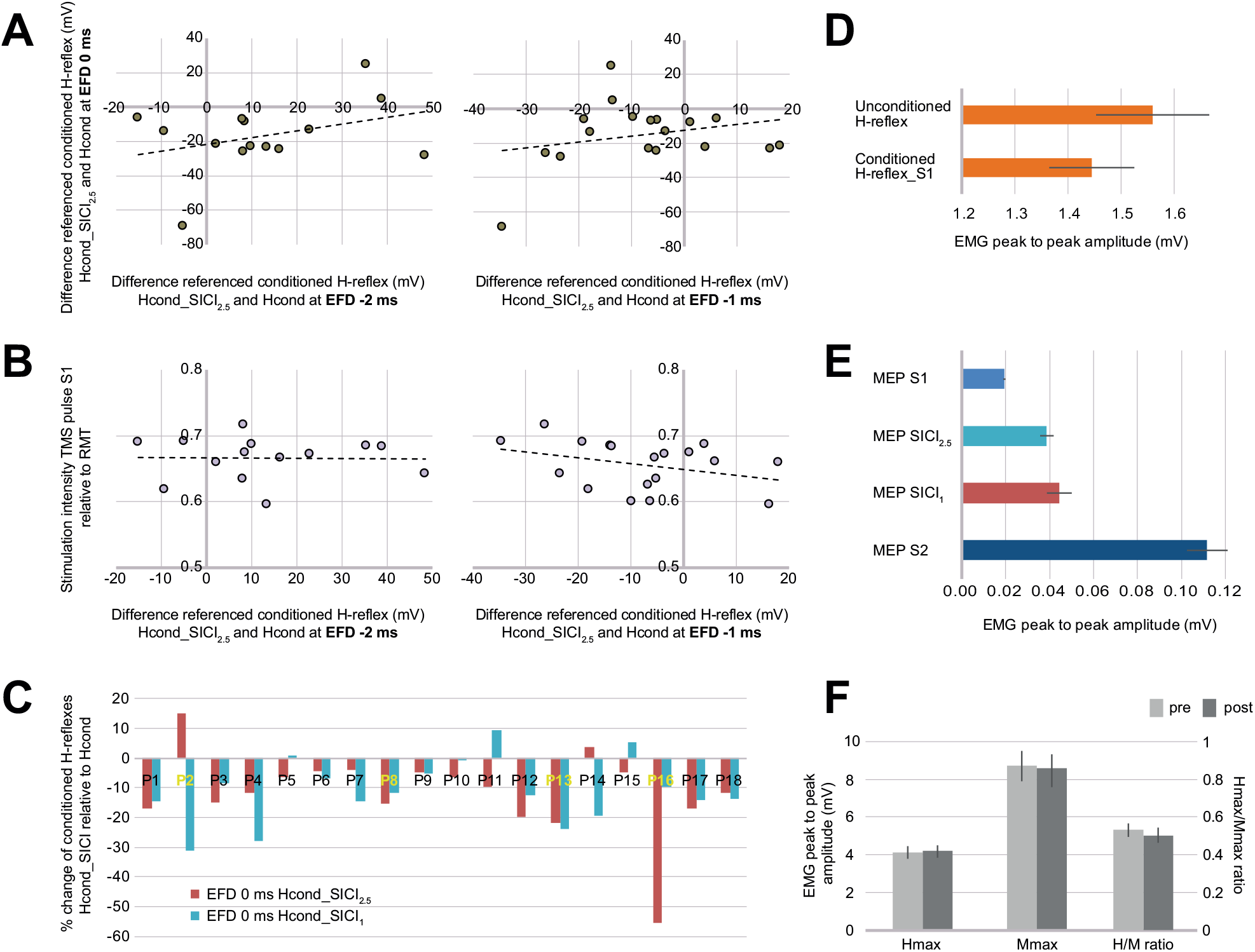
**A** depicts individual differences of referenced conditioned H-reflexes between Hcond_SICI_2.5_ and Hcond at EFD -2 ms (x-axis) and 0 ms (y-axis; left side) and at EFD -1 ms (x-axis) and 0 ms (y-axis, right side). The black dashed lines depict the regression of the data. There is no significant correlation between the displayed variables. **B** shows individual differences of referenced conditioned H-reflexes between Hcond_SICI_2.5_ and Hcond at EFD -2 ms (x-axis left side) and EFD -1 ms (x-axis right side) plotted against the stimulation intensity of the TMS pulse S1 relative to RMT (y-axis). The black dashed lines depict the regression of the data. There is no significant correlation between the displayed variables (left side: r = 0.34, P = 0.16; right side: r = 0.14, P = 0.63). **C** displays the individual percentual change of referenced conditioned H-reflexes at EFD 0 ms (early facilitation) in the Hcond_SICI conditions relative to the Hcond condition. Participants 2, 8, 13, and 16 are marked as they showed significant differences of referenced conditioned H-reflexes between the Hcond and Hcond_SICI_2.5_ conditions at EFD -2 ms or EFD -1 ms. P2 displayed a significant increase of referenced conditioned H-reflexes at EFD -2 ms for the Hcond_ SICI_2.5_ condition. P8, P13 and P16 showed a significant decrease of referenced conditioned H-reflexes at EFD -1 ms for the Hcond_ SICI_2.5_ condition. **D** displays the peak to peak amplitudes (mean ± SEM) of the unconditioned H-reflex and the H-reflex that was conditioned by S1 (Conditioned H-reflex_S1). Note that for conditioning we used a fixed ISI of +1 ms between PNS and the TMS pulse S1, which means that this ISI indicated different EFDs in different subjects, depending on the ISI of the individual early facilitation. **E** shows peak to peak amplitudes (mean ± SEM) for TMS compound potentials. Note that the SOL EMG background noise in our experiment was around 20 μV, thus S1 produced no visible SOL MEP in the EMG. **F** displays peak to peak amplitudes (mean ± SEM) for Hmax, Mmax and the H/M ratio obtained at the beginning and the end of each experiment.

## Discussion

Short interval intracortical inhibition (SICI) is well-described for the hand area of M1. A low intensity TMS pulse depresses corticospinal excitation produced by a subsequent suprathreshold test pulse. It is thought that local GABAAergic inhibitory connections are activated by the first stimulus and these depress the response to the second stimulus via local intracortical connections. We will refer below to this as a “SICI-like” effect.

We applied the same technique in the present experiments to the leg area innervating the SOL muscle. We had assumed that we would observe a similar phenomenon, and that a low intensity S1 would depress corticospinal output evoked by the suprathreshold S2 pulse. The data showed that these expected effects could be observed on the early, rapidly conducting outputs evoked by S2. Unexpectedly there were opposite effects on the later, slower conducting outputs evoked by the same low intensity pulse. As we argue below this implies that effects on the different outputs have fundamentally different mechanisms.

### Contrasting effects of the TMS pulse S1 on conditioned H-reflexes

Because the intensity of S2 was 1.2 RMT, it elicited an MEP in the SOL muscle. We could therefore confirm directly that, as expected by the SICI paradigm, this MEP was depressed to 60 % (Hcond_SICI_2.5_) and 65 % (Hcond_SICI_1_) of its mean amplitude. This is consistent with the SICI-like effect described above.

However, when we use MEPs to measure corticospinal excitation we assess the summation of all corticospinal neurons and connections that are influenced by TMS, which can be excitatory and inhibitory in nature. Further, slower conducting corticospinal volleys may not be equally represented in the MEP than faster conducting volleys based on the higher asynchronicity of their arrival at the spinal motoneurons. H-reflex conditioning accounts for these issues and measures the activity of corticospinal influences at the spinal motoneurons at multiple instants after TMS. At each of these instants, H-reflex conditioning can detect even weak descending corticospinal activity (Nielsen et al. 1993). This is possible as the afferent volley by PNS elevates the susceptibility of spinal motoneurons for synaptic input, making it easier for corticospinal descending activity to cross the firing threshold. Using this technique, we typically see that S2 evoked two periods of facilitation at the spinal level (see curve shape in the Hcond condition in Figure 2) (Taube et al. 2011). The first, lasting about 5 – 6 ms, correspond to corticospinal inputs that do very likely contribute to the MEP. The second, starting some 8 ms after the earliest arriving activity reflects later inputs to spinal motoneurons that may not be significantly reflected in the MEP.

The novel result in the present experiment is that an S1 which causes a “SICI-like” depression of the early period of facilitation, has the opposite effect (a pronounced facilitation) on later arriving inputs to the spinal cord. We can only speculate on the mechanisms underlying this later effect. Both S1 and S2 activate neurones in the cortex. If we assume that effects from these stimuli occur locally within stimulated areas of the cortex, the implication is that the cortical outputs modulated by S1 which arrive early at the segmental level differ from those that arrive later. For example, as effects at the early facilitation (EFD 0 ms) results from rapidly conducting corticospinal output from M1, the later facilitation may depend on indirect activation of slower conducting pathways from other motor areas like the premotor cortex. The present results could then indicate that S1 has opposite effects on outputs from M1 and premotor cortex. Another possibility is that S1 evokes output from M1 that influences subcortical relay pathways, for example in the reticular formation (Keizer and Kuypers 1984). It would then be possible for S1 to interact with cortical outputs evoked by S2 that travel via these relays (e.g. primary motor-premotor, primary motor-reticulospinal).

There was one feature of the effects of S1 which contrast previous results: it depressed the earliest arriving input to spinal motoneurons evoked by S2, referring to the depression of the referenced conditioned H-reflex at EFD 0 ms. This was true for both the 1 ms and 2.5 ms interval between S1 and S2. The effect at EFD 0 ms is consistent with the idea that the first-arriving input is not a D-wave, since D-waves are not depressed by SICI (Di Lazzaro, Oliviero, Profice, et al. 2001). In fact, the first arriving input from S1 is thought to be an I1-wave indicating that S1 trans-synaptically activates corticospinal neurones in the leg area as it does in the hand area of M1(Di Lazzaro, Oliviero, Profice, et al. 2001). However, in the hand area, SICI evoked by S1 has relatively little effect on the I1 inputs; most of its effect is on later I-inputs (usually I3) (Di Lazzaro, Restuccia, et al. (1998). The conflicting results between these findings and our study may be caused by the orientation of the induced current, which was in posterior-anterior direction (PA stimulation) in the present study. In the previous study of Di Lazzaro et al. (1998), SICI was produced with the magnetic coil positioned such that the current flow was in an anterior-posterior direction (AP stimulation). AP stimulation has been argued to produce largely I3 waves but less I1 waves, in contrast to stimulation where the current is PA, producing also consistent I1 waves (Di Lazzaro, Oliviero, Mazzone, et al. 2001). Thus underlying mechanisms of SICI with AP stimulation may be different than SICI with PA stimulation.

### S1 elicited corticospinal activity via fast conducting connections from M1

The earliest-arriving input evoked by S2 at the segmental level was aligned across individuals to EFD 0 ms. S2 had no effect on H-reflexes evoked prior to that time (EFD -2 and -1 ms). However, in the Hcond_SICI_2.5_ condition, there was a clear facilitation of the H-reflex at EFD - 2 ms. We propose that this was due to a small fast conducting corticospinal volley evoked by S1, which would reach the segmental level at the time when the afferent volley from PNS reaches the spinal motoneurons. Interestingly, at EFD -1 ms, Hcond_SICI_2.5_ depressed H-reflexes. Following the report of Cowan et al. (1986) of a similar effect in hand muscles, we suggest that this was caused by disynaptic inputs from spinal Ia inhibitory interneurons activated by descending input to the antagonist TA muscle. This inhibition cuts short the initial excitation evoked by S1, and depresses the H-reflex in SOL. Indeed, this period of depression, or at least of reduced facilitation is also evident in the Hcond curve of referenced conditioned H-reflex at EFD +1 ms. In this case the depression is caused by the suprathreshold S2 triggering presumably disynaptic inputs from spinal Ia inhibitory interneurons. Lack of overt inhibition in the Hcond_SICI_2.5_ condition may be due to the larger preceding initial facilitation at EFD 0 ms.

Very early H-reflex facilitation was not as clear in the Hcond_SICI_1_ condition. In this case, early descending activity induced by the fastest conducting corticospinal volley from S1 should arrive at the segmental level at EFD -1 ms. H-reflex testing at this time interval showed only a non-significant mean facilitation. We suggest that because S1 evokes only a small corticospinal volley, the fastest conducting corticospinal volley was not sufficiently strong to depolarize spinal motoneurons. In the case of Hcond_SICI_2.5_, the afferent volley by PNS depolarized spinal motoneurons 0.5 ms after the earliest possible arrival of excitation from corticospinal inputs, and thus later (0.5 ms) arriving volleys may be required to cross the firing threshold. Likely if we had tested Hcond_SICI_1_ at EFD -0.5 ms we might have seen an equally large facilitation of the H-reflex.

The strength of the modulation of referenced conditioned H-reflexes at EFD -2 ms and -1 ms with Hcond_SICI_2.5_ did not depend on the intensity of S1 relative to RMT. A stronger S1 pulse may produce stronger corticospinal activity and therefore a stronger modulation in the size of referenced conditioned H-reflexes. Lack of correlation may simply be the result of the limited range of S1 intensities. It might also reflect differences in the slope of the corticospinal input-output relationships between individuals: that is an intensity of e.g. 0.7 RMT may result in more or less corticospinal output in different individuals. The practical result is that an estimate of whether or not corticospinal activity is triggered by S1 cannot be based on the stimulation intensity alone. Instead, potential effects have to be measured.

Importantly, even though the TMS pulse S1 produced effects outside M1 and modulated referenced conditioned H-reflexes at EFD -2 ms and -1 ms, these effects were not related to the reduced conditioned H-reflexes seen at EFD 0 ms. The correlation analyses calculated for modulated referenced conditioned H-reflexes with Hcond_SICI_2.5_ between EFD -2 ms/-1 ms and EFD 0 ms yielded no significant results. Further support for this finding comes from the analyses of a reduced dataset. In these analyses, we discarded subjects that revealed a significant difference of referenced conditioned H-reflexes between conditions Hcond_SICI_2.5_ and Hcond at EFD -2 ms or -1 ms. These analyses showed no significant modulations of referenced conditioned H-reflexes at EFDs -2 ms and -1 ms, but still reduced referenced conditioned H-reflexes at EFD 0 ms with Hcond_SICI_2.5_. Thus at least for the fastest conducting corticospinal volley induced by S2 and probed at EFD 0 ms, the reduced excitation with Hcond_SICI_2.5_ is not mediated by corticospinal activity. The reduction is rather cortical in origin, highly likely because S1 activated a different subset of pronounced inhibitory synaptic inputs than the subsequent S2 that reduced output of corticomotoneuronal cells with fastest conducting corticospinal connections.

In 16 out of 18 tested subjects (89%) Hcond_SICI_2.5_ depressed referenced conditioned H-reflexes at EFD 0 ms. Of the two remaining subjects, one (P2) showed a larger facilitation of around 15%, and the second subject (P14) showed a rather small increase of around 4 %. The huge facilitation of 15 % in P2 (Figure 3C) is not consistent with the proposed cortical mechanism and can hardly be explained by natural variance of the physiological reponses. It is therefore noteworthy that P2 was the only subject we discarded from subsequent analyses of referenced conditioned H-reflexes because of significantly higher referenced conditioned H-reflexes at EFD -2 ms in Hcond_SICI_2.5_. We pointed out that the facilitation at EFD -2 ms is highly likely caused by descending input to spinal motoneurons from S1. Thus the unexpected facilitation seen at EFD 0 ms in P2 may be due to significant corticospinal activity from S1 obscuring depression at this time interval. In a similar manner, the extraordinary large reduction of referenced conditioned H-reflexes at EFD 0 ms with Hcond_SICI_2.5_ in P16 (Figure 3C) may be explained by a facilitated depression of SOL spinal motoneurons. This subject was discarded because the referenced conditioned H-reflexes at EFD -1 ms were significantly lower with Hcond_SICI_2.5_.

### Differences between Hcond_SICI_2.5_ and Hcond_SICI_1_

At EFD +1 ms, Hcond_SICI_1_ depressed the H-reflex more than Hcond, consistent with a SICI-like effect. However, there was no depression for Hcond_SICI_2.5_. At EFD +2 ms the situation was reversed. One possible explanation for this is that S1 evokes not only an early arriving corticospinal volley responsible for the effects observed at EFD -2 ms and -1 ms, but also a later volley arriving some 3 ms later. This would equate in the hand area to the I1 and I3 volleys that are often observed to be evoked by threshold TMS pulses (Di Lazzaro, Oliviero, et al. 1998). If so, then this facilitation would obscure the SICI-like depression at EFD +1 ms for Hcond_SICI_2.5_ and at EFD +2 ms for Hcond_SICI_1_.

### Transferability of the results to the upper limb

The results of the present study were obtained in the lower leg muscle SOL, and solid conclusions can therefore only be drawn for this muscle. We chose the lower limb as we obtained most of our experience with conditioned H-reflexes in the SOL, and as H-reflexes can be easily evoked in this leg muscle. Most SICI studies, however, investigated not the lower leg but hand and arm muscles of the upper limb. We do not make predictions of underlying neural mechanisms of SICI in the upper limb. Nevertheless, we presume that similar mechanisms may take place as in the leg, specifically with respect to the production of corticospinal activity by S1 and the contrasting effects of depression of faster corticospinal inputs to the spinal cord and facilitation of later arriving inputs.

### Effect of pairing S1 with PNS at later ISIs

In the first SICI study by Kujirai et al (1993) the authors conditioned an H-reflex in the upper limb with an S1 pulse at fixed ISIs of +1 ms to +10 ms and found no modulation in the size of the H-reflexes. These experiments were performed in three subjects. Indeed, for some ISIs S1 may not cause a modulation of the H-reflex. This is what we observed when we conditioned the H-reflex with the S1 pulse at a fixed ISI of +1 ms in our study, too. Without the coincidental finding at EFDs -2 ms and -1 ms with Hcond_SICI_2.5_ we would have likely concluded the same as Kujirai et al. (1993), namely that S1 caused no corticospinal activity. However, the negative result with fixed ISI(s) may simply be caused by overlapping excitatory and inhibitory effects from S1, also at the spinal level, cancelling each other out and producing no modulations of the H-reflex.

### Conclusion

In summary, in our study assessing the modulatory influence of subthreshold TMS pulse (S1) on the excitation of corticospinal volleys produced by a subsequent suprathreshold TMS pulse (S2), we saw mixed effects of depression and facilitation. Excitation of corticospinal volleys with faster conduction times was reduced, and excitation of corticospinal volleys with slower conducting times was increased. S1 produced corticospinal activity and not only local effects in M1 as it was previously assumed, and this activity may potentially mediate the changed excitation. At the level of M1, S1 very likely targeted different, preferentially inhibitory synaptic inputs to corticospinal output cells than S2, and this mechanism did very likely cause the reduced excitation of the fastest conducting corticospinal volley induced by S2. Based on these results, we believe that it is critically important to interpret underlying mechanisms of effects observed in studies applying SICI with caution.

## Grant

This study was supported by a research grant from the Deutsche Forschungsgemeinschaft (LE 2744/7-1).

